# A meta-analysis on uncertainty monitoring in four non-primate animal species: Pigeons, rats, large-billed crows, and bees

**DOI:** 10.1101/2020.12.03.411082

**Authors:** Zhizhen Qu, Sze Chai Kwok

## Abstract

Humans have the metacognitive capacity to be aware of what they do and do not know. While uncertainty monitoring has long been regarded as uniquely human, researchers in search of the polygenetic root of this ability have gathered evidence that primate species possess functional features parallel to humans. However, there were no systematic studies that quantitively take into account of extant data for these non-primate animals. Through a meta-analysis, we collected published data reported in 11 articles from 55 individual non-primate animals spanning over four species on the “opt-out” paradigm, the most prevailing paradigms used to test nonhuman animals’ uncertainty monitoring. We used chosen-forced advantage and opt-out rate to quantify animals’ performance results for computing the aggregated effect size for this literature. We found that these four NPA species process a significantly positive effect size for both scores and identified the moderators that have contributed to the inconsistencies across these studies. Implications for theories on metacognition are discussed.

## 1. Introduction

Humans can verbalize the mental experience of being uncertain, whereas animals do not have linguistic markers to communicate with us, preventing our direct access into understanding the basis of their mental state. To test animals’ metacognitive ability, a number of paradigms have been developed, such as uncertainty responses, betting, confidence ratings, and information-seeking tasks (Call & Carpenter, 2001; Middlebrooks & Sommer, 2011; Shields, Smith, Guttmannova, & Washburn, 2005). In the realm of non-primate animals (hereafter called NPAs), the paradigms are limited due to the difficulties involved in training them to understand such tasks. In the NPA literature, one of the most important tasks is the uncertainty morning “opt-out” task.

The first report of the opt-out task in a NPA species described a bottlenosed dophin (*Tursiops truncatus*) being able to “opt-out” when the decision threshold was at its perceptual limit in an auditory psychophysical task (Smith et al., 1995). It was argued that the dolphin’s opt-out behavior was an indicator that the dolphin knew when it felt uncertain. Similar experiments were conducted to test uncertainty monitoring abilities in rats, pigeons, large-billed crows, and bees. Researchers incorporated both easy and difficult trials either in a memory or perception task containing multiple difficulty levels and granted animals an “opt-out” option to let them decline to perform any given trials (for a comprehensive list of experiments, see **Table 2**).

**Table 1.** Moderator analysis results for chosen-forced advantage, opt-out rate, and composite score. The most significant moderator is highlighted in **bold**. The test order (prospective vs retrospective) is the most statistically significant moderator.

|  |  | Effect size and 95% confidence interval |  |  |  |  |  | Test of null (2-Tail) |  | Heterogeneity |  |  |
| --- | --- | --- | --- | --- | --- | --- | --- | --- | --- | --- | --- | --- |
|  |  | Number Studies | Point estimate | Standard error | Variance | Lower limit | Upper limit | Z-value | P-value | Q-value | df(Q) | P-value |
| Order |  |  |  |  |  |  |  |  |  | 2.607 | 1 | 0.106 |
|  | prospective | 4 | 0.025 | 0.028 | 0.001 | -0.028 | 0.079 | 0.925 | 0.355 |  |  |  |
|  | retrospective | 9 | 0.08 | 0.019 | 0 | 0.042 | 0.118 | 4.121 | 0 |  |  |  |
| Species |  |  |  |  |  |  |  |  |  | 5.537 | 3 | 0.136 |
|  | bee | 1 | 0.081 | 0.058 | 0.003 | -0.032 | 0.194 | 1.401 | 0.161 |  |  |  |
|  | large-billed crow | 1 | 0.01 | 0.052 | 0.003 | -0.092 | 0.112 | 0.193 | 0.847 |  |  |  |
|  | pigeon | 4 | 0.025 | 0.027 | 0.001 | -0.028 | 0.078 | 0.929 | 0.353 |  |  |  |
|  | rat | 5 | 0.104 | 0.026 | 0.001 | 0.052 | 0.155 | 3.959 | 0 |  |  |  |
| Domain |  |  |  |  |  |  |  |  |  | **6.163** | 1 | 0.013 |
|  | memory | 4 | 0.018 | 0.021 | 0 | -0.024 | 0.06 | 0.853 | 0.394 |  |  |  |
|  | perception | 7 | 0.088 | 0.018 | 0 | 0.052 | 0.125 | 4.789 | 0 |  |  |  |
|  |  | Number Studies | Point estimate | Standard error | Variance | Lower limit | Upper limit | Z-value | P-value | Q-value | df(Q) | P-value |
| Order |  |  |  |  |  |  |  |  |  | **4.655** | 1 | 0.031 |
|  | prospective | 4 | 0.087 | 0.087 | 0.008 | -0.083 | 0.257 | 1.008 | 0.313 |  |  |  |
|  | retrospective | 8 | 0.327 | 0.069 | 0.005 | 0.191 | 0.462 | 4.727 | 0 |  |  |  |
| Species |  |  |  |  |  |  |  |  |  | 0.155 | 3 | 0.984 |
|  | bee | 1 | 0.131 | 0.652 | 0.426 | -1.148 | 1.41 | 0.201 | 0.841 |  |  |  |
|  | large-billed crow | 1 | 0.38 | 0.281 | 0.079 | -0.172 | 0.932 | 1.35 | 0.177 |  |  |  |
|  | pigeon | 4 | 0.295 | 0.116 | 0.014 | 0.067 | 0.523 | 2.539 | 0.011 |  |  |  |
|  | rat | 4 | 0.29 | 0.113 | 0.013 | 0.069 | 0.512 | 2.569 | 0.01 |  |  |  |
| Domain |  |  |  |  |  |  |  |  |  | 0.033 | 1 | 0.856 |
|  | memory | 3 | 0.317 | 0.137 | 0.019 | 0.05 | 0.585 | 2.325 | 0.02 |  |  |  |
|  | perception | 7 | 0.288 | 0.092 | 0.008 | 0.108 | 0.468 | 3.131 | 0.002 |  |  |  |
|  |  | Number Studies | Point estimate | Standard error | Variance | Lower limit | Upper limit | Z-value | P-value | Q-value | df(Q) | P-value |
| Order |  |  |  |  |  |  |  |  |  | **2.562** | 1 | 0.109 |
|  | prospective | 4 | 0.092 | 0.051 | 0.003 | -0.008 | 0.191 | 1.809 | 0.07 |  |  |  |
|  | retrospective | 9 | 0.189 | 0.034 | 0.001 | 0.123 | 0.255 | 5.632 | 0 |  |  |  |
| Species |  |  |  |  |  |  |  |  |  | 0.382 | 3 | 0.944 |
|  | bee | 1 | 0.11 | 0.129 | 0.017 | -0.142 | 0.362 | 0.854 | 0.393 |  |  |  |
|  | large-billed crow | 1 | 0.19 | 0.147 | 0.022 | -0.099 | 0.479 | 1.289 | 0.197 |  |  |  |
|  | pigeon | 4 | 0.162 | 0.064 | 0.004 | 0.037 | 0.288 | 2.533 | 0.011 |  |  |  |
|  | rat | 5 | 0.191 | 0.057 | 0.003 | 0.08 | 0.302 | 3.367 | 0.001 |  |  |  |
| Domain |  |  |  |  |  |  |  |  |  | 0.675 | 1 | 0.411 |
|  | memory | 4 | 0.114 | 0.043 | 0.002 | 0.059 | 0.229 | 3.311 | 0.001 |  |  |  |
|  | perception | 7 | 0.189 | 0.033 | 0.001 | 0.123 | 0.254 | 5.656 | 0 |  |  |  |

**Table 2.** 15 studies on uncertainty monitoring paradigm. Studies that are included in this meta-analysis is denoted with an asterisk *.

| **NO.** | **Author** | **Year** | **Journal** | **Species** | **No. of subjects** |
| --- | --- | --- | --- | --- | --- |
| 1 | Smith and Schull | 1989 | unpublished data | Brown rat | 6 |
| 2 | Teller | 1989 | unpublished undergraduate thesis | Rock pigeon | 6 |
| 3 | Smith et al | 1995 | Journal of Experimental Psychology: General | Common bottlenose dolphin | 1 |
| 4 | Inman and Shettleworth* | 1999 | Journal of Experimental Psychology: Animal Learning and Cognition | Rock pigeon | 4 |
| 5 | Sole et al* | 2003 | Psychonomic Bulletin & Review | Rock pigeon | 3 |
| 6 | Foote and Crystal* | 2007 | Current Biology | Brown rat | 3 |
| 7 | Sutton and Shettleworth* | 2008 | Journal of Experimental Psychology: Animal Behavior Processes | Rock pigeon | 7 |
| 8 | Adams and Santi* | 2011 | Learning & Behavior | Rock pigeon | 7 |
| 9 | Angel* | 2010 | PhD thesis | Brown rat | 2 |
| 10 | Nakamura et al | 2011a | Animal Cognition | Rock pigeon | 6 |
| 11 | Nakamura et al | 2011b | Animal Cognition | Bantam | 3 |
| 12 | Goto and Watanabe* | 2012 | Animal Cognition | Large-billed crow | 3 |
| 13 | Perry and Barron* | 2013 | PNAS | Honeybee | 10 |
| 14 | Templer et al* | 2017 | Animal Cognition | Brown rat | 9 |
| 15 | Yuki and Okanoya* | 2017 | Journal of Experimental Psychology: Animal Learning and Cognition | Brown rat | 4 |
| 16 | Yuma Osako et al* | 2018 | Scientific Reports | Brown rat | 3 |

In this paradigm, two key parameters measuring task success are the chosen-forced advantage and the opt-out rate (Smith, Shields, & Washburn, 2003). These two parameters is a metric first reported in Teller (1989) and Inman and Shettleworth’s (1999) studies. The prediction is that an animal should perform better in chosen trials with an “opt-out” option compared with “forced” ones wherein an option to decline the test is absent. During the former condition, the animal should only accept trials that it considers itself capable of responding correctly. Under a metacognitive account assumption, the “opt-out” rate should thus be higher in trials with higher difficulty.

In the past three decades, 11 articles reported performance in four different NPA species; yet, different studies yield different task results. The considerable inconsistencies might be attributed to differences in how these studies were conducted. A meta-analysis approach is the ideal method for understanding NPA animals’ group performance when the findings are discrepant across studies. Meta-analyses not only provide a general and combined estimate of NPA’s performance in the opt-out task, but also help to identify potential moderators that may have contributed to the inconsistencies in the findings. To the best of our knowledge, to date, there have been no systematic reviews or metaanalyses exploring NPA’s performance on this metacognitive task, nor the potential moderators contributed to the inconsistencies across different studies.

In this study, we collected data from opt-out studies that used the chosen-forced advantage and the opt-out rate to quantify animals’ performance from 11 published articles. We excluded two species from the analysis because one species (dolphin) just has data point for one single animal subject, and the other (bantam) did not participate in a standardized paradigm. We also computed a composite score, which is the average of an individual animal’s chosen-forced advantage and the opt-out rate to offer us a 2-in-1 measure for evaluating its performance. Our aim was to (1) estimate the overall effect size for the chosen-forced advantage, opt-out rate, and composite score across four different NPA species; and (2) understand how the factors species, task order (prospective vs retrospective), and task domain (memory vs perception) would influence these animals’ performances.

## 2. Method

### 2.1 Selection of studies

We followed the conventional procedure for the systematic reviews and metaanalyses (PRISMA) approach for the data collection. We first conducted a literature search for papers that contained empirical studies on NPA performance on the opt-out task prior to October 2020 through PubMed, PsycNET, Web of Science, and ProQuest Dissertations & Theses Global. Search terms included (‘metacognition’) and (‘animals’ or ‘non-primate’). We also manually searched for papers that cited the most relevant reviews (Robert R Hampton, 2009; Smith, 2009; Smith et al., 2003) in the field. Articles were included in the meta-analysis if they met the following criteria: (a) written in English; (b) published in journals or dissertations; (c) using a standard opt-out paradigm and reported both chosen-forced advantage and opt-out rate. In cases wherein the chosen-forced advantage and the opt-out rate were not available in the articles, we have reached out to relevant authors and obtained the raw data. In cases wherein we were not able to obtain the original data, we used estimated values based on figures published in the papers. Some studies used a variant of the paradigm, or conducted multiply experimental sessions aiming to understand how advanced these animals’ meta-ability is (e.g., test on an immediate transfer to new stimuli). We will discuss these variants in the discussion section, and these studies/sessions were not included into the standard metaanalysis; and (d) only NPA species were included as subjects. We however excluded two unpublished studies (one on pigeons and one on rats), and a study conducted on a dolphin because the study was conducted on one single subject, giving us insufficient sample size to run the analysis. In the end, we included 11 papers covering pigeons, rats, large-billed crows, and bees in the meta-analysis.

### 2.2 Coding of studies

The following information was extracted from each of the included studies: *author(s) and publication year; type of publication* (journal or dissertation); *species; test domain* (perception or memory); *test order* (prospective or retrospective); *sample size; chosen-forced advantage; opt-out rate; data status* (collected or estimated).

### 2.3 Calculating effect size

To allow cross-species and cross-experiment comparisons, we evaluated all of the extant data using the metric composed of chosen-forced advantage and opt-out rate. For each individual animal, the chosen-forced advantage is calculated by obtaining the average “percentage correct on chosen trials” minus “percentage correct on forced trials” on all difficulty levels. The opt-out rate is calculated as “percentage of tests declined at the highest difficulty level” minus “lowest difficulty level”. We expected the chosen-forced advantage to be statistically higher than zero if NPAs were able to use the opt-out option when they feel uncertain. The composite score is the average of an individual animal’s chosen-forced advantage and the opt-out rate. We extracted the average rate for these three scores, as well as the standard error of the means among the individual animals within-study for the meta-analysis.

### 2.3 Data analysis

The Comprehensive Meta-Analysis software version 2 (Borenstein, 2005) was used to synthesize the data and perform the statistical analysis. Due to the heterogeneity in sampling methods, assessment instruments, and sample size across studies, a randomeffects model was used to estimate the effect size for the three scores. Cochran’s Q and the I^2^ statistic was used to assess the degree of heterogeneity across included studies. Subgroup analyses were used to examine the sources of heterogeneity and the key moderators that contributed to the heterogeneity. Publication bias was evaluated with the funnel plots and Egger’s test.

## 3. Results

From the eleven studies that we obtained the three scores, the pooled effect size for chosen-forced advantage, opt-out rate, and composite score are 0.060 (95% CI: 0.028 to 0.092, I^2^=91.823%), and 0.297 (95% CI: 0.154 to 0.440, I^2^=93.957%), and 0.173 (95% CI: 0.112 to 0.233, I^2^=94.458%) respectively, indicating a small but significant effect size. We also visualized the distribution of all the data in Figure 3, showing that most animals have positive scores in both measurements.

**Figure 1.**
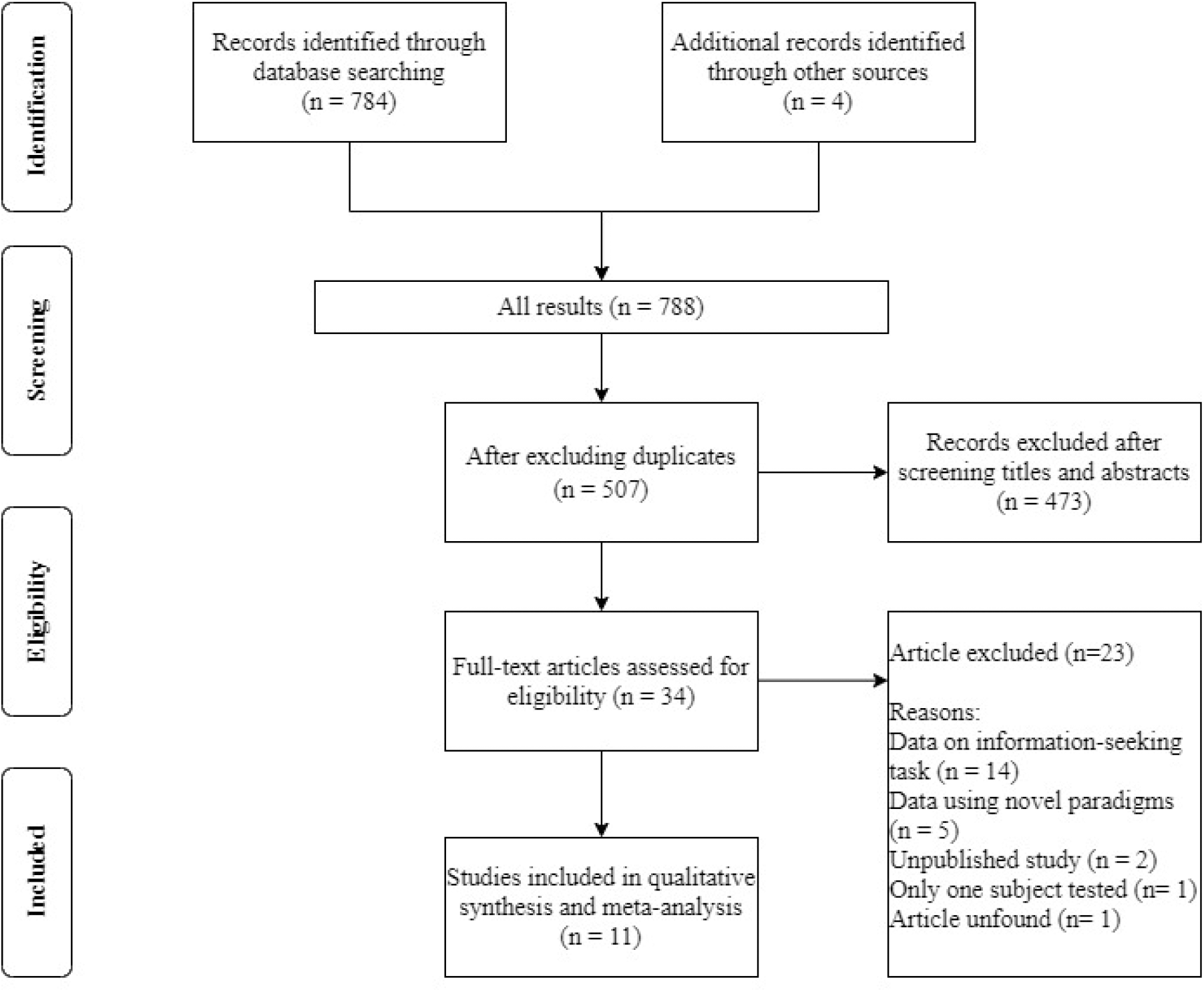
Study selection flowchart.

**Figure 2.**
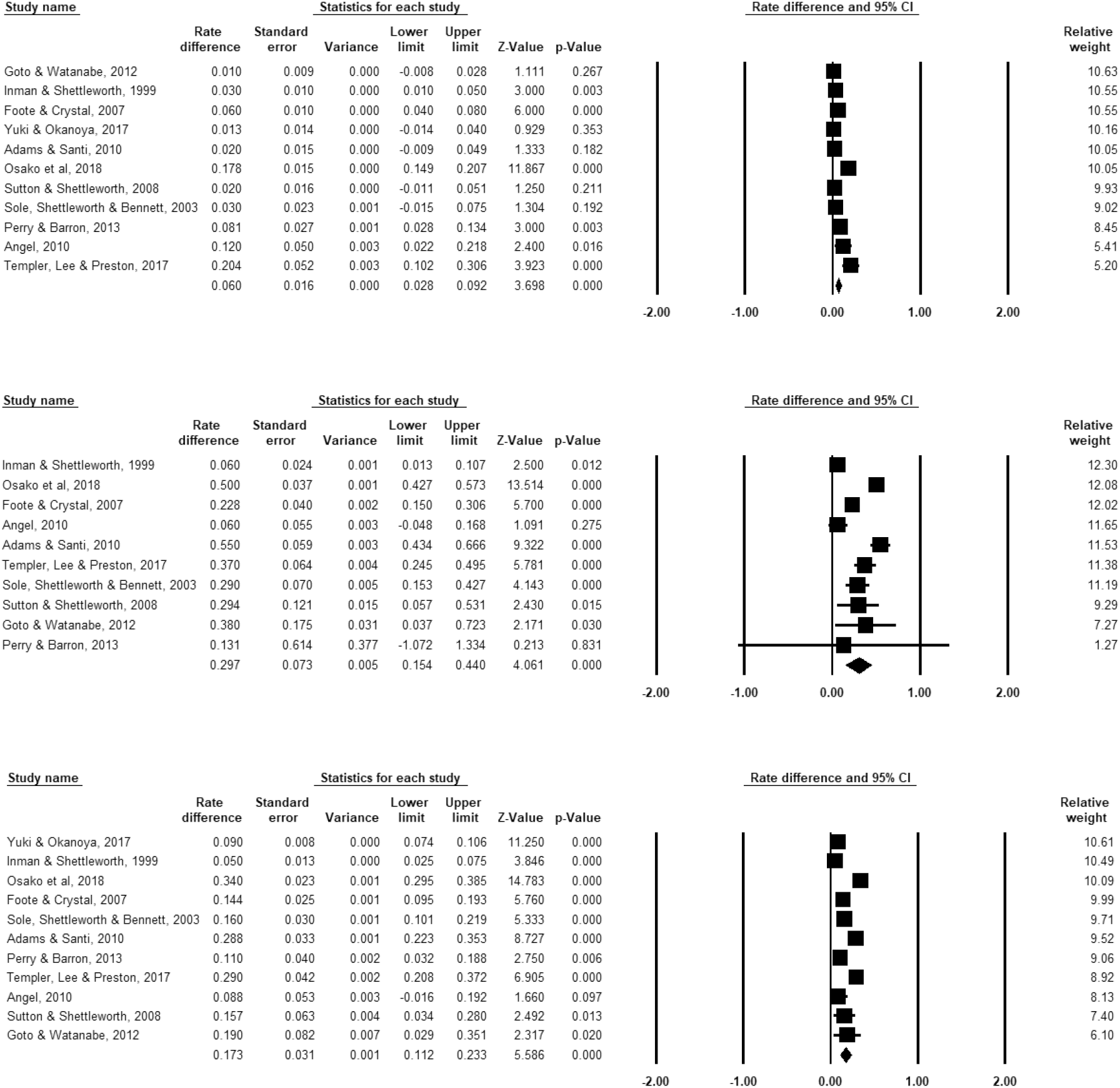
Forest plot for chosen-forced advantage (top panel), opt-out rate (middle panel), and composite score (bottom panel). An effect size is estimated for each study, and the relative weight represents (rightest column) how much this specific study contributed to the overall analysis. Horizontal lines show 95% confidence interval; the diamond at the bottom in each panel represents the point estimate and confidence interval of the pooled effect size. Note that for Yuki & Okanoya (2017), we only found the mean opt-out rate for the 4 rats participated in the experiment, thus we exclude this study from the forest plot for the opt-out rate (middle panel), because we were not able to acquire the standard error of the means among the individual animals.

**Figure 3.**
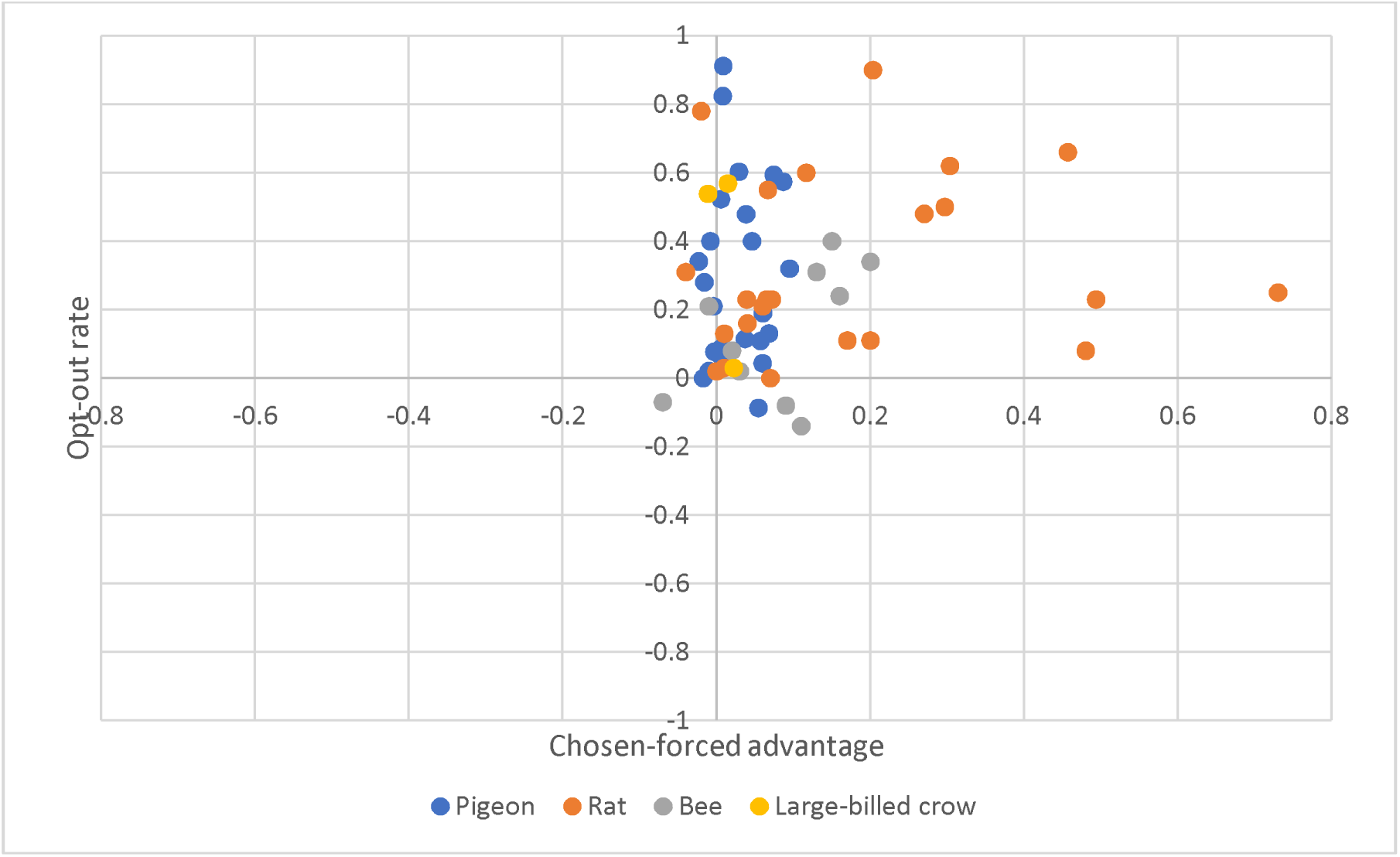
Scatter plot for chosen-forced advantage and opt-out rate. Each dot represents an individual animal. The colors denote the type of species.

Substantial heterogeneity was also identified, with Q(10)□=122.291 (*p*□ <□ .001) for chosen-forced advantage; Q(9)□=148.942 (*p*□<□ .001) for opt-out rate, and Q(10)□= 180.455 (*p*□<□ .001) for composite score. A limitation of Cochran’s Q test is that it might be underpowered when few studies have been included or when event rates are low, thus we tested the I^2^, which provides an estimate of the percentage of variability in results across studies that is due to real differences and not due to chance. We found I^2^ 90.559% for chosen-forced advantage, I^2^□=96.934% for opt-out rate, and I^2^□=94.859 for composite score. According to Higgins et al (2003), I^2^□values of 25 %, 50 %, and 75 % indicate low, moderate and high heterogeneity, respectively. Our data contain very high heterogeneity.

## 5. Moderator analysis

To delve further into the factors that lead to the high heterogeneity, we conducted subgroup analysis for the three scores. Results showed that domain contributed significantly to the heterogeneity for the chosen-forced advantage (Q=6.163, *p*=0.013), test order contributed significantly to the heterogeneity for opt-out rate (Q=4.655, *p*=0.031) and for the composite score (Q=2.562, *p*=0.109). Averaging the Q values for the three scores, test order (prospective vs retrospective) proved to be the most statistically significant moderator (**Table 1**).

## 6. Publication bias

To examine if publication bias existed in the study, we used funnel plots for chosen-forced advantage, opt-out rate, and the composite score; a symmetric plot would indicate a lack of publication bias (Sterne, Becker, & Egger, 2005). Egger’s tests were also performed to examine whether the assumption of a symmetrical distribution of effect sizes is viable; a *p* value greater than .05 would suggest a lack of sufficient evidence for publication bias (Egger, Smith, Schneider, & Minder, 1997). Egger’s tests revealed that there was no publication bias for chosen-forced advantage (t=1.246, *p* =0.244; Figure 4 top panel), opt-out rate (t=1.034, *p* =0.331; Figure 4 middle panel) and the composite score (t=1.859, *p* =0.096; Figure 4 bottom panel).

**Figure 4.**
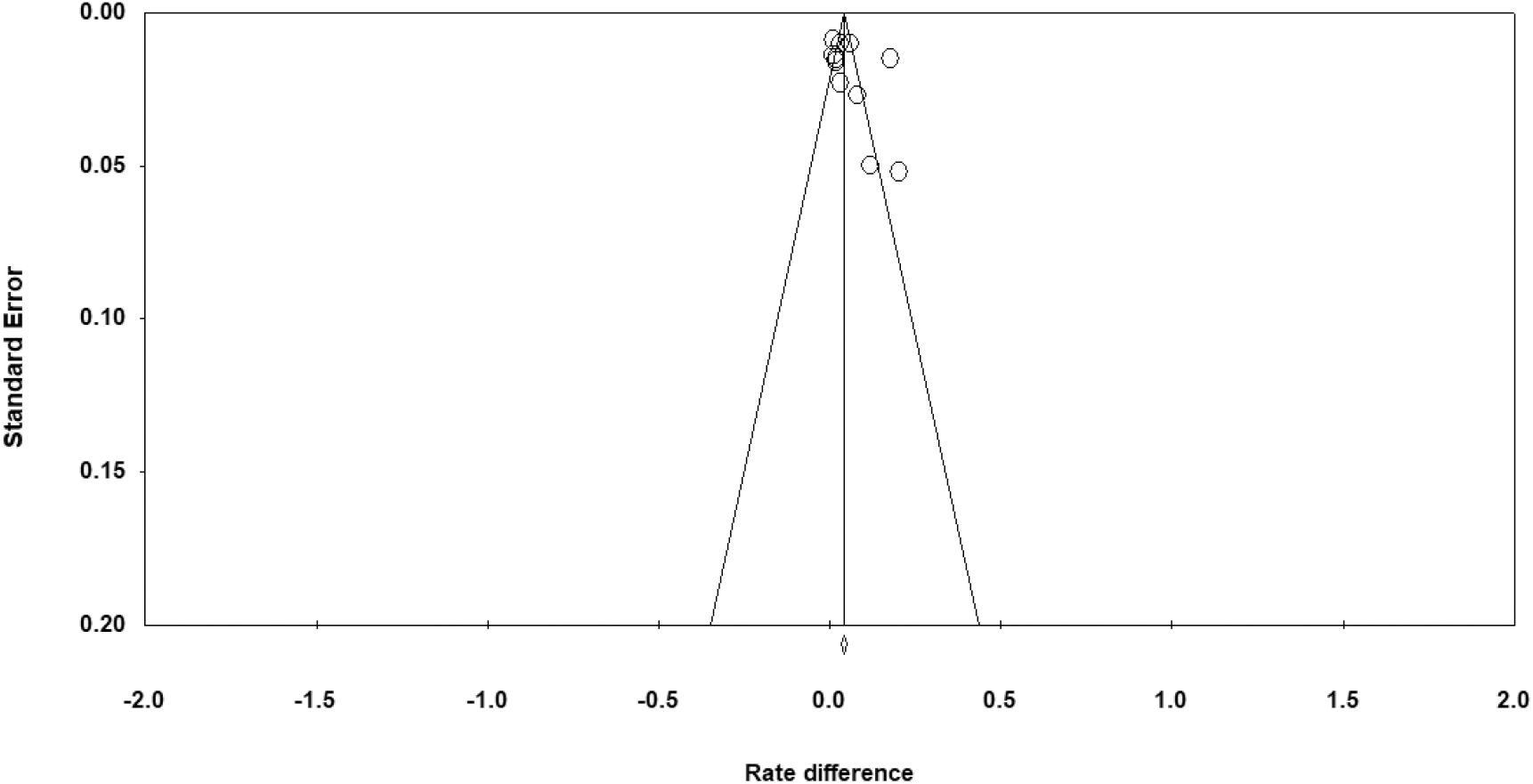
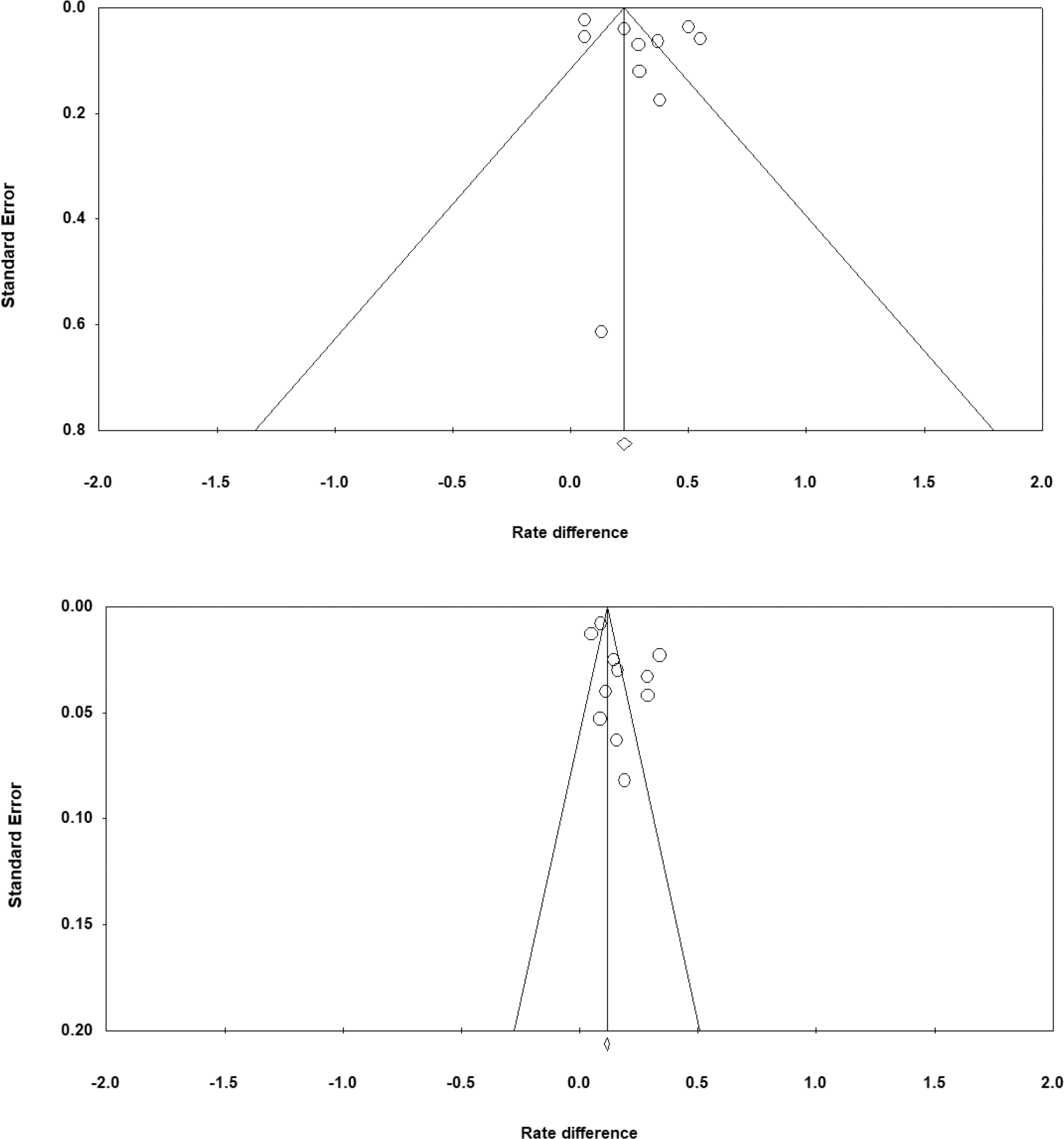
Funnel plots for chosen-forced advantage (top panel), opt-out rate (middle panel), and composite score (bottom panel).

## 7. Discussion

Our findings that these four NPA species passed the uncertainty monitoring test at the group level carry theoretical implications for the field. Different levels of metacognition have been proposed by scholars to describe animals’ meta behaviors, such as object versus meta level (Shields et al., 2005), first-order versus second-order (Crystal & Foote, 2009), public mechanism versus private mechanism (R. R. Hampton, 2009). The meta-like performance on this task remains debatable as the measures do not unambiguously reflect second-order (i.e., metacognitive) computations. To address this issue, a number of transfer tasks have been developed. In these transfer task, animals were tested on whether they could transfer the concept of opting out to a novel task. The impetus for developing such tasks is to show if animals are able to pass a transfer task following a primary task, which would indicate that their uncertainty monitoring ability in a task-independent cognitive state (Washburn, Smith, & Shields, 2006).

Moreover, researchers have tested if uncertainty monitoring ability is transferable to novel stimuli in pigeons (Nakamura, Watanabe, Betsuyaku, & Fujita, 2011; Sole, Shettleworth, & Bennett, 2003), bantams (Nakamura et al., 2011), and bees (Perry & Barron, 2013). Five out of six pigeons and two out of three bantams in Nakamura’s study (2011) generalized their uncertainty responses to novel stimuli at least once. Four of the ten bees could transfer the concept of opting out to a novel task (Perry & Barron, 2013). In these tasks, subjects demonstrated a generalized mechanism whereby a first-order (cognitive) representation is internally assessed through a second-order (metacognitive) process that directly evaluates its quality, although it remains unknown if such a mechanism contains introspection. Such performance has prompted theorists to land a “middle ground” between the low level and the high level to discuss animals’ metabehavior (Metcalfe & Son, 2012; Smith, Couchman, & Beran, 2012). It opens up possibilities to interpret the existing NPA metacognition literature in relation to their evolutionary significance.

In this meta-analysis, we determined that test order played an important role in leading to the heterogeneity across studies. The result is in alignment with a number of test rests on non-primate and primate that animals more often showed retrospective memory than prospective memory (Morgan, Kornell, Kornblum, & Terrace, 2014), and the argument that metacognitive judgments are less accurate given prospectively than retrospectively (Siedlecka, Paulewicz, & Wierzchoń, 2016). Interestingly, we did not find the factor species to be a determinant factor leading to the heterogeneity of animals’ performance, and this is in alignment with the emerging evidence that metacognitive-like abilities detected from animals from various distant families. One study on pharaoh’s ants found that individual ants spontaneously upregulate or downregulate pheromones depending on the reliability of their own memories (Czaczkes & Heinze, 2015). This demonstrates certain metamemory-like ability in ants, echoing other insect studies wherein honeybees were found to be able to monitor their uncertainty (Perry & Barron, 2013). It is possible that the evolution of metacognition is not a representation of a linear sequence of cortex-dependent evolution, but rather as representatives of different clades that diverged at different points during evolution. We also found that test domain was not a decisive factor in the heterogeneity of NPAs’ performances. In primate and human literature, it remains controversial whether metacognition shows domain specificity, with supporting evidence for both domain-specific and domain-general metacognitive representations (Morales, Lau, & Fleming, 2018; Ye, Zou, Lau, Hu, & Kwok, 2018). Since these NPA species do not possess the cortex that can be specialized in either one of the tasks, thus it is likely that they are using a domain general strategy in handling such tasks.

## 8. Conclusion

We present a cross-species comparative meta-analysis on four species of NPA’s meta-ability measured by the opt-out paradigm. By aggregating the existing evidence in support of some degree of meta-ability in these animals, we showed that the four NPA species pass the uncertainty monitoring test. We identified test order as the principal factor that contributed the most to the heterogeneity of these animals’ performance. We hope this study would help consolidate the research in the field and stimulate research towards a less anthropocentric direction in the study of metacognition.

## Acknowledgement

SCK was supported by the National Natural Science Foundation of China (Grant No. 32071060), the Science and Technology Commission of Shanghai Municipality (Grant No. 201409002800) and JORISS project grants.

The authors declared no potential conflicts of interest with respect to the research, authorship and/or publication of this article.

